# Identifying Crohn’s disease signal from variome analysis

**DOI:** 10.1101/216432

**Authors:** Yanran Wang, Yuri Astrakhan, Britt-Sabina Petersen, Stefan Schreiber, Andre Franke, Yana Bromberg

## Abstract

**Background:** After many years of concentrated research efforts, the exact cause of Crohn’s disease remains unknown. Its accurate diagnosis, however, helps in management and even preventing the onset of disease. Genome-wide association studies have identified 140 loci associated with CD, but these carry very small log odds ratios and are uninformative for diagnoses.

**Results:** Here we describe a machine learning method – AVA,Dx (Analysis of Variation for Association with Disease) – that uses whole exome sequencing data to make predictions of CD status. Using the person-specific variation in these genes from a panel of only 111 individuals, we built disease-prediction models informative of previously undiscovered disease genes. In this panel, our models differentiate CD patients from healthy controls with 71% precision and 73% recall at the default cutoff. By additionally accounting for batch effects, we are also able to predict individual CD status for previously unseen individuals from a separate CD study (84% precision, 73% recall).

**Conclusions:** Larger training panels and additional features, including regulatory variants and environmental factors, e.g. human-associated microbiota, are expected to improve model performance. However, current results already position AVA,Dx as both an effective method for highlighting pathogenesis pathways and as a simple Crohn’s disease risk analysis tool, which can improve clinical diagnostic time and accuracy.

## Background

Crohn’s disease (CD) is a chronic inflammatory bowel disease (IBD) of the gastrointestinal tract with an incidence between 0.1–16 per 100,000 persons worldwide[1]. Chronic inflammation may occur in any part of the gastrointestinal tract and may in some cases also manifest extraintestinally. A combination of genetic and environmental factors is involved in disease etiology, with genetics accounting for 75%[2]. Genome-wide association studies (GWAS) contributed to the understanding of the genetic architecture of CD and have, so far, identified 140 significantly associated loci[3]. These results elucidate the underlying molecular disease pathways, contributing to the understanding of the fundamental biology behind CD pathogenesis. GWAS results highlight the roles of ER stress, barrier integrity, innate immunity, autophagy, cytokine production, lymphocyte activation, the response to bacteria and specifically the role of the JAK-STAT-pathway. With few exceptions, individual risk loci confer only a modest effect on disease susceptibility. Taken together the known loci explain approximately 13% of disease[3]. Thus, CD diagnosis still requires exploration of the patients’ clinical history, a physical exam, blood tests (mainly for detecting inflammation markers and excluding infections as a cause) and, as the gold standard, an invasive endoscopy and pathology reports for visual confirmation of the inflammation. Several serologic markers, primarily anti-*Saccharomyces cerevisiae* antibody (ASCA) and perinuclear anti-neutrophilic cytoplasmic antibody (pANCA), have recently been suggested to be clinically useful for diagnosis. However, these markers are not accurate enough to precisely diagnose Crohn's disease on their own and are, therefore, considered an addition to conventional tests. Moreover, for six[4] to nine[5] percent of IBD patients the diagnosis changes during the course of disease, suggesting that some patients are erroneously diagnosed and in some cases even not treated for the right disease.

The predictive value of genetic testing for the disease-associated variants is controversial since the identified mutations generally exhibit weak correlation and do not identify causative patterns. Still, computational predictions, based on 30 GWAS CD loci, have attained a fairly high accuracy[6, 7] with ROC AUC (area under the receiver operating characteristic curve) of 0.71 that can be further improved to 0.74 by incorporating family history. In another study, a logistic regression model attainted even higher reported predictive performance (ROC AUC = 0.86) by training on 573 GWAS loci in over 13,300 individuals[8].

While CD GWAS-based models may have high predictive ability, they require large panel sizes for identification of significant loci and for model building. Whole exome or genome data can provide an alternative, pathogenesis pathway-oriented, perspective, as it has been suggested that over 96% of the predicted functionally significant human exome variants are rare or private single nucleotide variants (SNVs)[7]. Also note that using the exome-available markers in our data set, the above-described GWAS-based regression model performed worse than expected (ROC AUC = 0.63 for our CD-train cohort).

Here we show that health status predictions based on functional effects of all individual-specific non-synonymous variants can be used to discriminate between patients (CD) and healthy individuals (HC). Interestingly, a model built using our variant and gene scoring rules for the known GWAS locus genes[3] was unable to outperform random guessing. However, by selecting from the GWAS set of genes a subset of those that the Pascal[9] method (for computing gene and pathway scores from SNP-phenotype association summary statistics) identifies as being most relevant to CD, we improved model performance to ROC AUC of 0.70, similar to findings reported earlier[6]. Note that model performance was far worse when our scoring function for these genes only accounted for the load of variants per gene (variant burden), rather than their effects on molecular functionality. These results suggest that changes to molecular functions of affected genes are more representative of disease-associated pathway deficiencies than the number of variants per pathway alone.

We used computational feature selection techniques to identify a CD-relevant gene set from our exome data, increasing model performance even further (mean ROC AUC = 0.74). We termed the combination of our gene selection and model training approach AVA,Dx -- Analysis of Variation for Association with Disease X; *i.e.* we believe that AVA,Dx is generic enough to be applied to other diseases. Note that this approach did not incorporate any prior knowledge of CD biology and our selected genes were not significantly overlapping with any of the previously identified sets of genes. These findings suggest that our approach highlighted new, likely causative, genes that are part of Crohn’s pathogenesis pathways.

Further, to test the true predictive performance of our model, we corrected for batch effects individually for every person from another CD/HC panel, and applied AVA,Dx. Remarkably, our method achieved 84% precision and 73% recall at the default cutoff.

Finally, we note that our approach has so far required only a very small set of people to draw conclusions. Moreover, we only included the exonic information from Whole Exome Sequencing (WES) data, while there actually is a lot of regulatory information in WES data as well[10]. Larger training panels and additional features, including regulatory variants and, potentially, environmental factors (*e.g.* human-associated microbiota), are expected to improve model performance. However, current results already position AVA,Dx as both an effective method for highlighting pathogenesis pathways and as a simple CD-risk analysis tool, which can improve clinical diagnostic time and accuracy.

## Results

### AVA,Dx pipeline

We constructed the AVA,Dx pipeline as outlined in Fig. 1 (a more detailed pipeline in Additional file 1: Figure 1) and Methods. Briefly, we cleaned up the data, performed predictions of functional effects of variants[11, 12] and converted the latter into per-gene scores of individual-specific functionality. To build predictive models we considered externally determined disease genes (e.g. GWAS and literature-identified genes) and extracted our own disease-gene sets via computational feature selection (FS). Note that in this manner we identified previously unreported CD-related genes. Support Vector Machine[13] (SVM) models using these gene sets were tested in *leave-one-out* cross-validation on the training data and on (batch effect corrected) previously unseen data. In permutation testing, we showed that our prediction performance was significantly non-random.

**Figure 1.**
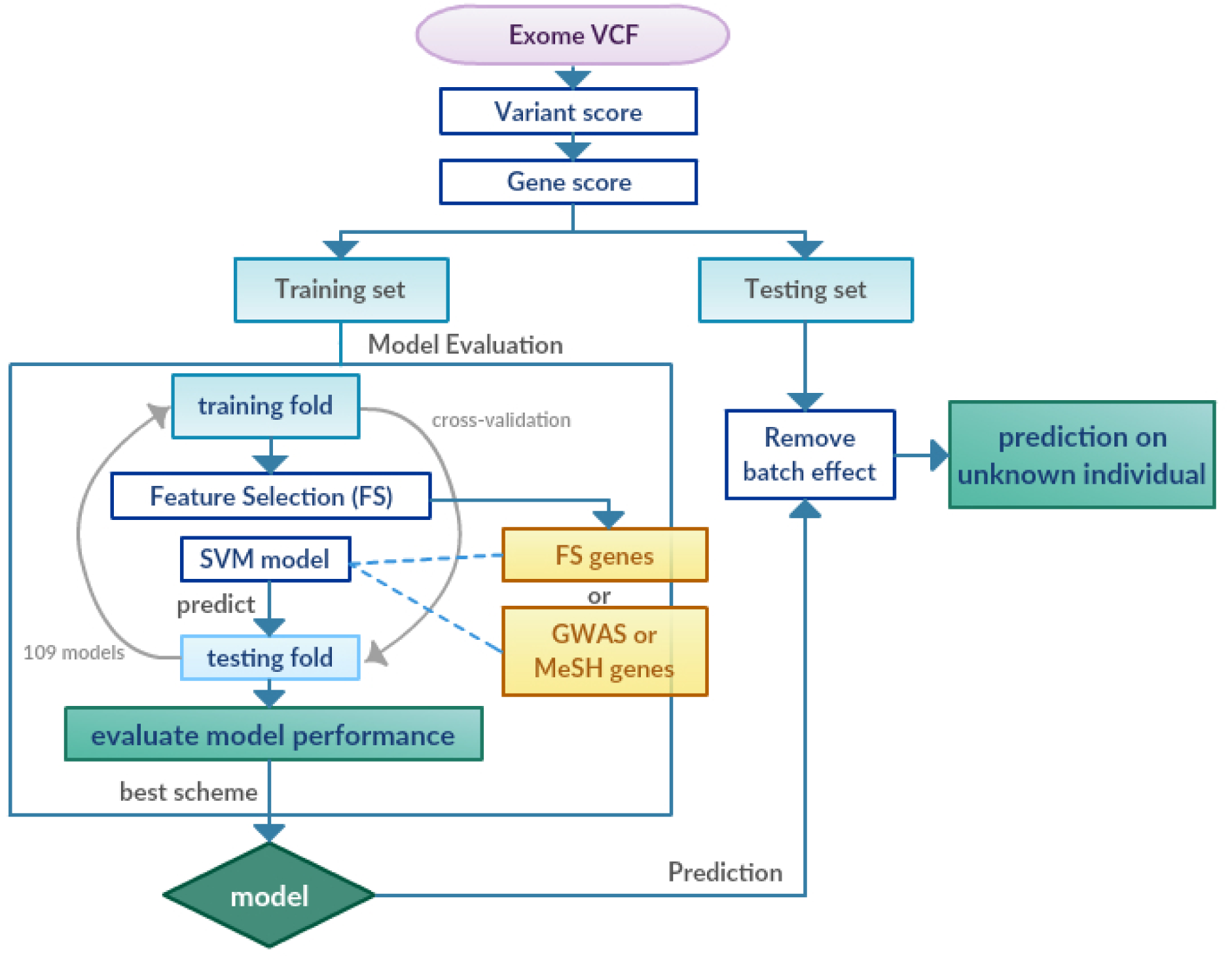
AVA,Dx pipeline summary. A simplified pipeline of AVA,Dx (Analysis of Variation for Association with Disease). Input data is in Variant Call Format (VCF) file from Whole Exome Sequencing. Different versions of *gene_scores* are calculated as described in Methods. Models for prediction of Crohn’s disease status are evaluated by cross-validation. Best *gene_score* scheme, FS algorithm, and, finally, SVM model, are selected on the basis of performance in cross-validation. These are further used for prediction of unknown individuals. A more detailed flowchart is in Additional file 1: Figure 1.

### Data Cleanup: Missing calls contribute to variant burden differences between CDs and HCs

Even after removing all sex chromosome variants and variants predicted to be false positives[14] (Methods), the variant burdens of CD and HC individuals were still significantly different (Additional file 1: Figure 2A, p-val < 0.005, two-sided Kolmogorov-Smirnov, KS, test[15], Additional file 2 and 3) in both CD-train and CD-test panels. In evaluating this observation, we found that this difference was largely due to the difference in missing variant calls. There are two ways to address missing data[16, 17]: remove individuals with missing calls in a minimum number of variants or remove variant loci missing from a certain number of individuals. In an effort to retain as many individuals for our study as possible, we chose the latter approach. We conservatively filtered out all variants that had a missing genotype in even one individual; *i.e.* after filtering, all individuals in our study had genotypes for all the same variant loci. In this step, we removed 25,393 (12.8% of 198,406 in total) variant loci in the CD-train panel and 19,005 (12.8% of 147,998 in total) in the CD-test panel. After filtering, the transition/transversion ratio has increased from 2.61 to 2.69 and from 2.62 to 2.70 in CD-train and CD-test, respectively, in line with accepted metrics of ~3.0 for exome[14]. On average, 16,661 ± 125 and 16,516 ± 157 (mean ± s.d.) exonic variants per individual sample remained in the CD-train and CD-test panels, respectively. There were no variant burden differences between CDs and HCs in both panels after filtering (p-val = 0.9 and 0.4 for CD-train and CD-test panel, respectively, Additional file 1: Figure 2B, Additional file 4 and 5). We used only the filtered data sets for all further analyses.

### Association tests fail to identify CD-associated SNVs

We performed Fischer exact association testing[17] with Benjamini-Hochberg correction[18] on the clean CD-train panel. Two variants were significant (rs782662688 and rs3088113, corrected p-val = 0.03149 and 0.04159, MAF = 0.0046 and 0.2587 from ExAC[19], respectively, Additional file 1: Figure 3 and Additional File 6). These two variants are in coding regions of SHANK2 and CKAP4 genes, respectively. Interestingly, these two are both only found in the healthy individuals, suggesting a possible protective role in CD.

### Known CD genes are uninformative of individual health status

To identify disease signal, for each gene we calculated scores that (1) give different weights to different variant types, based on the severity of their effects on protein function[11, 12] (*gene_score*). Additionally, (2) *easy_gene_score* that assigns a fixed value for all types of exonic variants, (3) *binary_gene_score* that uses binary effect/no-effect predictions for non-synonymous variants, (4) exonic variant burden (*EV_gene_score*), and (5) non-synonymous variant burden (*NSV_gene_score*) versions of gene scoring were also evaluated. We built SVM models in *leave-one-out* cross-validation on the CD-train panel, using randomly chosen subsets of various sizes of all human genes (*ALL set*), as well as the genes mapping to the known GWAS-established CD loci[3] (*GWAS set*), literature-annotated CD genes (*MeSH set*), and Swiss-Prot annotated CD genes (*SP set*). All model building procedures and data sets are described in detail in the Methods section.

As expected, models that were built on the *ALL set* genes were similar in performance to random guessing with any *gene_score* scheme. Increasing the number of genes per subset used for model building did not improve performance (Additional file 1: Figure 4 to 8). Interestingly, *GWAS* and *MeSH* sets had similar results as the *ALL* set (Fig. 2A, Additional file 1: Figure 4 to 8). The best performance was achieved by using all 18 *SP* genes (*SP*_*max*_) with *NSV_gene_score* (ROC AUC = 0.55 and area under precision-recall curve, PR AUC = 0.66, Additional file 1: Figure 8). While disappointing, the inability of these models to differentiate healthy individuals from disease affected ones, suggests that there is no sequencing or scoring artifact that could differentiate CDs and HCs in CD-train panel.

**Figure 2.**
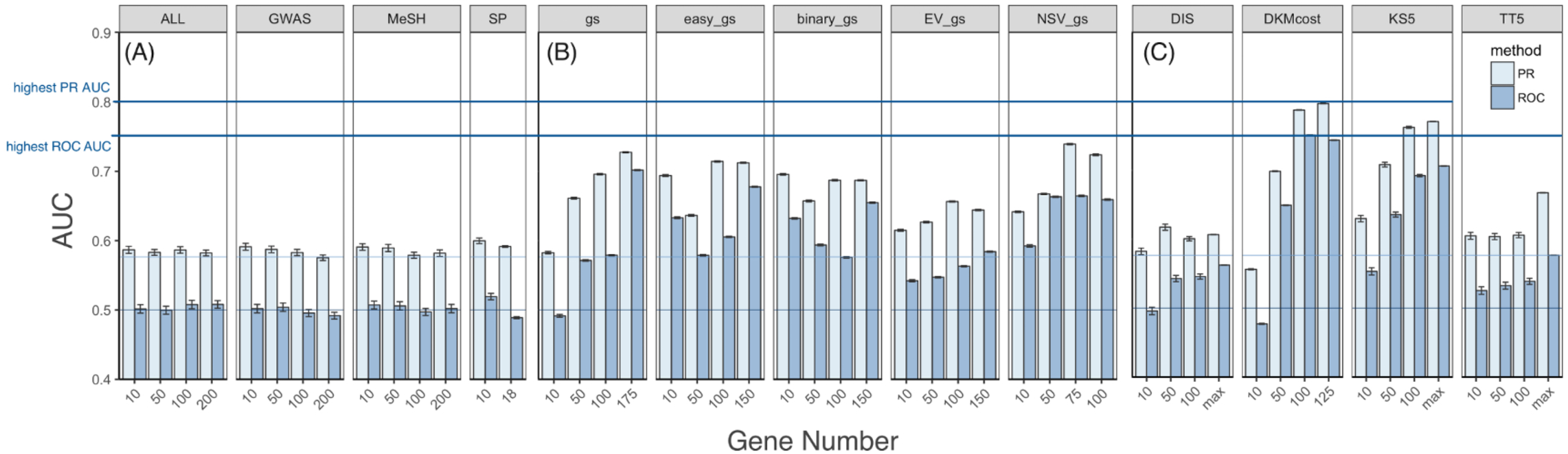
Performance of FS genes achieves highest value than other gene sets. Bar height indicates the AUC for precision/recall (PR, light blue) and ROC (dark blue) curves in 109-fold leave one out cross-validation on the CD-train panel (Methods). SVM models were re-trained 100 times, using **(A)** different numbers of randomly selected genes from *ALL*, *GWAS*, *MeSH*, and *SP* sets and *gene_score* (Methods) scoring **(B)** different numbers of top ranked *PascalGWAS* genes using different scoring schemes (Methods), **(C)** different numbers of top ranked *DKMcost* genes and randomly selected *DIS*, *KS5*, and *TT5* genes (Methods) by gene_score. Note that the number of genes used in each iteration was the same (e.g. top ranked 10 genes), but the resampling of training fold individuals was different. Error bars are standard errors over 100 iterations. The dark blue line at AUC=0.5 represents random for ROC AUC, while the light blue line at AUC=0.58 represents random for PR AUC. For simplicity, only 10, 50, 100, and the best performing gene numbers are shown here, and full results are in Additional file 1: Figures 4 to 12.

### Pascal-ranked CD genes significantly differentiate CDs from HCs

Since the *GWAS set* has 925 genes (Additional file 7), a random sample of these does not necessarily represent the true GWAS power. Instead, we took the *Pascal*-ranked[9] CD GWAS genes (*PascalGWAS set*, Methods, Additional file 8) and built SVM models for *leave-one-out* cross-validation using top ranked subsets of genes in various sizes. The *Pascal*-ranked genes achieved much better performance than other external sets (Fig. 2B). Using *gene_score* our models achieved the highest ROC AUC of 0.70 (PR AUC = 0.73, with *PascalGWAS*_*175*_ genes, *i.e.* top-ranked 175 genes from *PascalGWAS set*). By further permuting the CD/HC labels in cross-validation (Methods), we empirically showed that the performance of our models is significantly non-random (ROC/PR permutation p-val = 0.001/0.011). The *NSV_gene_score* had slightly worse but still significant performance (ROC/PR AUC = 0.66/0.74, achieved with *PascalGWAS*_*75*_ genes, permutation p-val = 0.013/0.001). The other three scoring schemes (*easy_gene_score*, *binary_gene_score* and *EV_gene_score*) had better performance with *Pascal*-ranked genes than with the other external sets, but they did not perform as well as *gene_score* and *NSV_gene_score* (Additional file 1: Figure 4-8). Thus, from here on we only considered the *gene_score* and the *NSV_gene_score* for all model building. Note that here and in all fixed gene sets, the only source of the difference in model performance is the differential resampling of the training individuals of the minor class (Methods).

### FS genes outperform top-ranked Pascal genes in differentiating CDs from HCs

We further evaluated the performance of the computationally extracted FS genes (*DIS/DISO*, *KS5*, *TT5* and *DKMcost sets;* Methods). Since both *gene_score* and *NSV_gene_score* performed relatively well compared with other scoring scheme, we used them both for the FS analyses. Trivially, the best performance (ROC/PR AUC = 1/1) was achieved by the *DISO* (*Disease Overfitted*) *sets* of more than 100 genes, defined as genes that were not affected in any of the CD-train HCs (Methods; Additional file 1: Figure 9 and 10).

*KS5*_*max*_ (all genes in *KS5 set*, ROC/PR AUC = 0.71/0.77, permutation p-val = 0.033/0.022; Additional file 1: Figure 11 and 13, Additional file 9) and *DKMcost*_*125*_ (top-ranked 125 genes from *DKMcost sets*, ROC/PR AUC = 0.74/0.80, permutation p-val = 0.014/0.010; Additional file 1: Figure 11 and 14, Additional file 10) *sets* using *gene_score* both outperformed *PascalGWAS*_*175*_ (Fig. 2B and 2C). For the *KS5 set*, including more genes slightly improved performance; *i.e.* performance of *KS5*_*r10*_ (random 10 genes from *KS5 set*) < *KS5*_*r25*_ < … < *KS5*_*max*_ (all genes in *KS5 set* were used, Additional file 1: Figure 11). This was not the case for *DKMcost* whose performance had reached a peak at 125 genes before dropping off. Note, however, that the maximum number of *KS* genes was 127, suggesting that models may simply not benefit from additional genes. *TT5*_*max*_ and *DIS*_*max*_ genes also had outperformed random guessing, but were not as good as *KS5*_*max*_ or *DKMcost*_*125*_ genes (Fig. 2C). Also note that the FS sets were never overfitted to the data, as feature selection was performed in a *leave-one-out* fashion, *i.e.* excluding the testing individual. Thus, our results suggest that FS selected genes can differentiate CDs from HCs in our data, particularly using signal that is not available to the statistics-based methods like GWAS.

The *KS5* genes *NSV_gene_score* selected genes (28-41 genes per iteration) failed to differentiate CDs from HCs in CD-train (ROC/PR AUC = 0.32/0.47). The *DKMcost* set was better, but also performed poorly (ROC/PR AUC = 0.65/0.73; Additional file 1: Figure 12). These results suggested that *gene_score* is more informative than *NSV_gene_score,* highlighting the importance of using the severity of functional effects of variants in evaluating disease genetics.

### Feature selection identifies known and previously undescribed CD genes

In the *leave-one-out* cross-validation, *PascalGWAS*_*175*_, *DKMcost*_*125*_ *and KS5*_*max*_ *sets* performed well in differentiating CDs from HCs in the CD-train panel (Fig. 2). Interestingly, the *KS5*_*max*_ and *DKMcost*_*125*_ *sets* only overlapped with *PascalGWAS*_*175*_ genes by one to five and two to four genes (average p-val = 0.34 and 0.25, hypergeometric test[20], background of *ALL* genes, Table 1), respectively. On the other hand, the *KS5*_*max*_ set significantly overlapped with the *DKMcost*_*125*_ genes (57 to 69 genes; average p-val = 6.64e-93). We also found that the FS sets (*DKMcost*_*125*_ *and KS5*_*max*_) did not significantly overlap with most of the external sets (*GWAS*, *MeSH*, and *PascalGWAS*_*175*_), but all external sets overlapped with each other (all with p-val < 0.05, Table 1 and Additional file 11). Note as an exception that while *Swiss-Prot* had no overlap with *KS5*_*max*_, the overlap of the former with *DKMcost*_*125*_ was significant (NOD2, MDR1, and DMBT1 genes of only 18 in *Swiss-Prot*), highlighting well-known CD genes extracted computationally without prior knowledge.

**Table 1.** **Gene set overlap summary**. Grey cells on the diagonal are the gene numbers in each set, e.g. *GWAS set* has 925 genes. The upper triangle describes the numbers of overlapped genes between two sets. The lower triangle shows the p-value of the corresponding overlap calculated by hypergeometric test in the background of *ALL* genes. Highlighted cells indicate significant overlap. Since there are 109 FS sets (*DKMcost*_*125*_ and *KS5*_*max*_) for each FS method from 109-fold cross-validations, italic values are used to indicate the mean overlap numbers (ranges in the parenthesis).

|  | GWAS | MeSH | Swiss-Prot | Pascal<br>GWAS<sub>175</sub> | DKM<br>cost<sub>125</sub> | KS5<sub>max</sub> |
| --- | --- | --- | --- | --- | --- | --- |
| **GWAS** | 925 | 203 | 9 | 121 | 12<br>(10 to 13) | 7<br>(5 to 9) |
| **MeSH** | 7.91E-15 | 1824 | 18 | 66 | 18<br>(14 to 21) | 16<br>(12 to 20) |
| **Swiss-Prot** | 6.69E-07 | 1.15E-16 | 18 | 8 | 2<br>(2 to 3) | 0<br>(0 to 1) |
| **PascalGWAS<sub>175</sub>** | 2.68E-102 | 1.15E-16 | 2.05E-11 | 175 | 3<br>(2 to 4) | 2<br>(1 to 5) |
| **DKMcost<sub>125</sub>** | 0.162 | 0.346 | 0.011 | 0.249 | 125 | 63<br>(57 to 69) |
| **KS5<sub>max</sub>** | 0.729 | 0.567 | 0.579 | 0.344 | 6.64E-93 | 113 |

Our FS techniques identified some known (*GWAS* and *MeSH*) genes. For example, both *DKMcost*_*125*_ and *KS5*_*max*_ contained the CD-associated LRRK2[21] and the uncharacterized KIAA1109 genes that also appeared in GWAS and MeSH sets. Additionally, *DKMcost*_*125*_ genes NOD2, LSP1, and CCR6 and *KS5*_*max*_ genes IL19 and ATF4 also appeared in *GWAS* and *MeSH* and were incidentally identified in follow-up studies as possibly CD-causative[22–26]. Overall, however, few genes appeared both in the FS sets and in the experimentally derived sets. The performance of the *KS5*_*max*_ and *DKMcost*_*125*_ models thus suggests that computational FS methods identify previously unsuspected CD genes as well.

### CD relevant genes interact

We used gene set overrepresentation analysis[27] to check if *DKMcost*_*125*_ genes selected from the entire CD-train are enriched in known molecular pathways (Methods). We also compared the enriched pathways of these genes with those of the *KS5*_*max*_ and the *PascalGWAS*_*175*_ genes (Additional file 12). FS found several significantly enriched pathways that were not identified by *PascalGWAS*_*175*_, but likely related to CD, *e.g.* antimicrobial peptides[28–31], apoptosis-related pathways[32, 33], cGMP effects[34–36], neutrophil degranulation[37, 38], innate immune system[39, 40], *etc.* Additionally, the protein-protein interaction network[27] of *DKMcost*_*125*_ genes (Fig. 3) suggested additional genes/proteins, which were not found by FS but may be relevant to CD; *e.g* the TAF1 and the HNF4A transcription factors regulate many *DKMcost*_*125*_ genes, including the infamous NOD2[22]. HNF4A was annotated as CD-associated in previous studies[41–43]. TAF1, on the other hand, needs further evaluation, but preliminary analysis shows that it contains a bromodomain, which may be critical in inflammation in general and bowel inflammation specifically in CD[44, 45].

**Figure 3.**
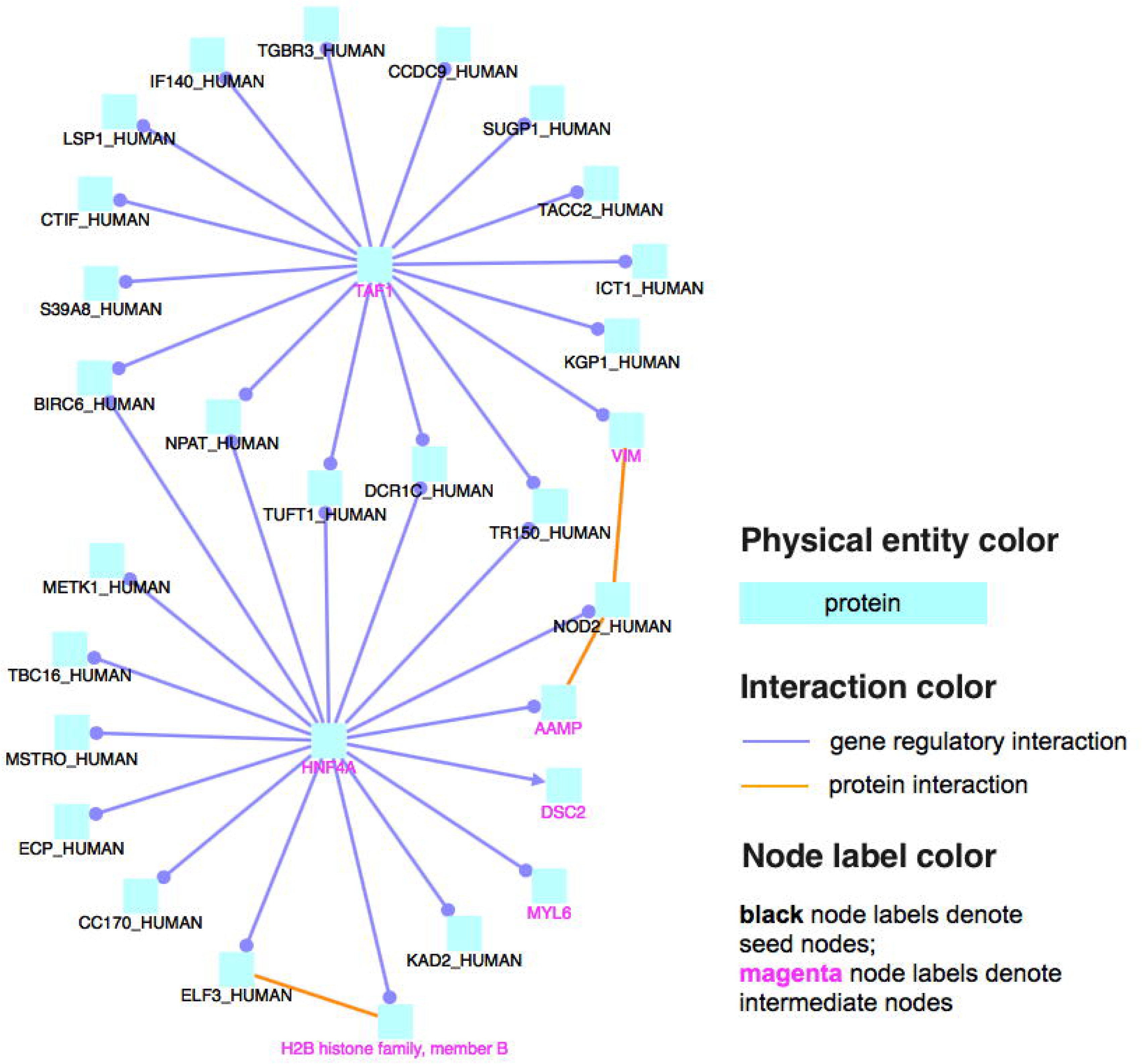
Induced Network of FS genes. *DKMcost*_*125*_ genes were the input for induced network analysis to ConsensusPathDB. Induced network analysis finds interacting genes/proteins (orange: protein interaction; blue: genetic interaction), starting from the input list (black titles node titles) and adding in others in the database (pink title nodes). Only high-confidence interactions are shown. TAF1 and HNF4 transcription factors are connected to genes/proteins found in the *DKMcost*_*125*_ set, indicating that they may be crucial to CD.

### Different pathogenesis pathways in early and late-onset CD

Earlier studies[46–49] highlighted a significant genetic difference between early onset and late onset of CD. To test this, we divided the CD-train panel into CD-train_early_ and CD-train_late_ according to the onset age of CD individuals (Methods); note that both panels contained the same 47 HC individuals. We performed feature selection on CD-train_early_ and CD-train_late_ panels separately and evaluated the performance of models trained with the selected genes (Additional file 1: Figure 15 and 16). The top-ranked *DKMcost* 275 and 200 genes from CD-train_early_ and CD-train_late_ (Additional file 13), respectively, were used for gene pathway overrepresentation analysis. In addition to some of the same pathways identified for the entire CD-train panel, *DKMcost*_*275*_ genes from CD-train_early_ were significantly enriched in endosomal/vacuolar pathway, antigen presentation, hedgehog pathway, graft-versus-host disease, type 1 diabetes mellitus, proteasome degradation, *etc*. (Additional file 14). However, no known molecular pathway was significantly enriched in *DKMcost*_*200*_ genes from CD-train_late_. These results suggest that genetically encoded pathways of CD pathogenesis are more obvious in the early-onset patients, while late onset CD may be more environmentally (or microbially) driven – a theory that we will test with more data in the future.

### Batch effect across panels

While filtering removed the overall variant count discrepancies between HCs and CDs within a panel, differences between panels remained. Regardless of health status, individuals from CD-train had significantly more variants than those from CD-test (Additional file 15 and 16). Furthermore, there were differences in variant burdens by variant type; *e.g*. CD-train had significantly more synonymous and heterozygous non-synonymous SNVs and less stop-losses and heterozygous stop-gains than CD-test. In total, CD-train and CD-test shared 98,795 SNVs, while, additionally, CD-train had 74,218 (42.9%) and CD-test had 30,198 (23.4%) unique SNVs. This “batch effect” may indicate, among other issues, a difference in coverage of sequence regions of different sequencing platforms[50, 51].

To evaluate the severity of the batch effect, we combined the CD-train and CD-test individual *ALL gene_score* profiles into one panel and performed clustering using *pam*[52]. Individuals clustered precisely according to batch (Additional file 1: Figure 17), indicating that obvious batch differences that would be likely to negatively impact prediction. To proceed with testing our CD prediction models on previously unseen data we had to first remove the batch effect. We first evaluated using the ComBat[53] algorithm on the combined panel of CD-train and CD-test (Methods). After adjustment of the *gene_score* value by ComBat, we were no longer able to separate batches (35.87% agreement between cluster members and batch labels, Additional file 1: Figure 18). Note that the originally selected *DKMcost*_*125*_ genes still maintained their ability to differentiate CD-train CDs from HCs in *leave-one-out* cross-validation after ComBat adjustment (ROC/PR AUC = 0.74/0.80, Additional file 1: Figure 19).

### Clinical application of AVA, Dx

To apply our method in a real-life situation, where one individual is evaluated for CD from his/her exome sequencing data, we built *PascalGWAS*_*175*_ *and* the best-performing *DKMcost*_*125*_ gene-based prediction models using the entire CD-train panel. Note that *DKMcost*_*125*_ genes selected here and were not identical to the ones used in cross-validation (>77% overlap). For prediction of each individual from the CD-test panel, we applied ComBat[53] to adjust *gene_scores* of the entire CD-train panel and the one individual, respectively (Methods).

To select the default cutoff in AVA,Dx score for calling an individual healthy or CD-affected we plotted the prediction scores for each individual from CD-train in cross-validation and selected a cutoff that best differentiated CDs from HCs (Additional file 1: Figure 20). The default cutoff was thus set at 14.3 (Methods) for all future evaluations. Note that in cross-validation, this meant that 47 of 64 CD patients were correctly identified, as were 28 of 47 healthy controls – 71% precision and 73% recall (F-measure = 0.72, Matthews correlation coefficient, MCC = 0.33).

In predicting the health status of all CD-test individuals the *PascalGWAS*_*175*_ model was nearly random (ROC/PR AUC of 0.56/0.84). Our *DKMcost*_*125*_ model, however, reached ROC/PR AUC = 0.68/0.88 (permutation p-val = 0.041/0.035) in predicting the health status of all individuals from CD-test (Additional file 1: Figure 21). That is, at default cutoff, using this model we were able to correctly identify 37 of 51 CD patients and 8 of 15 healthy controls (Fig. 4, precision=84%, recall=73%, F-measure=0.78, MCC=0.23).

**Figure 4.**
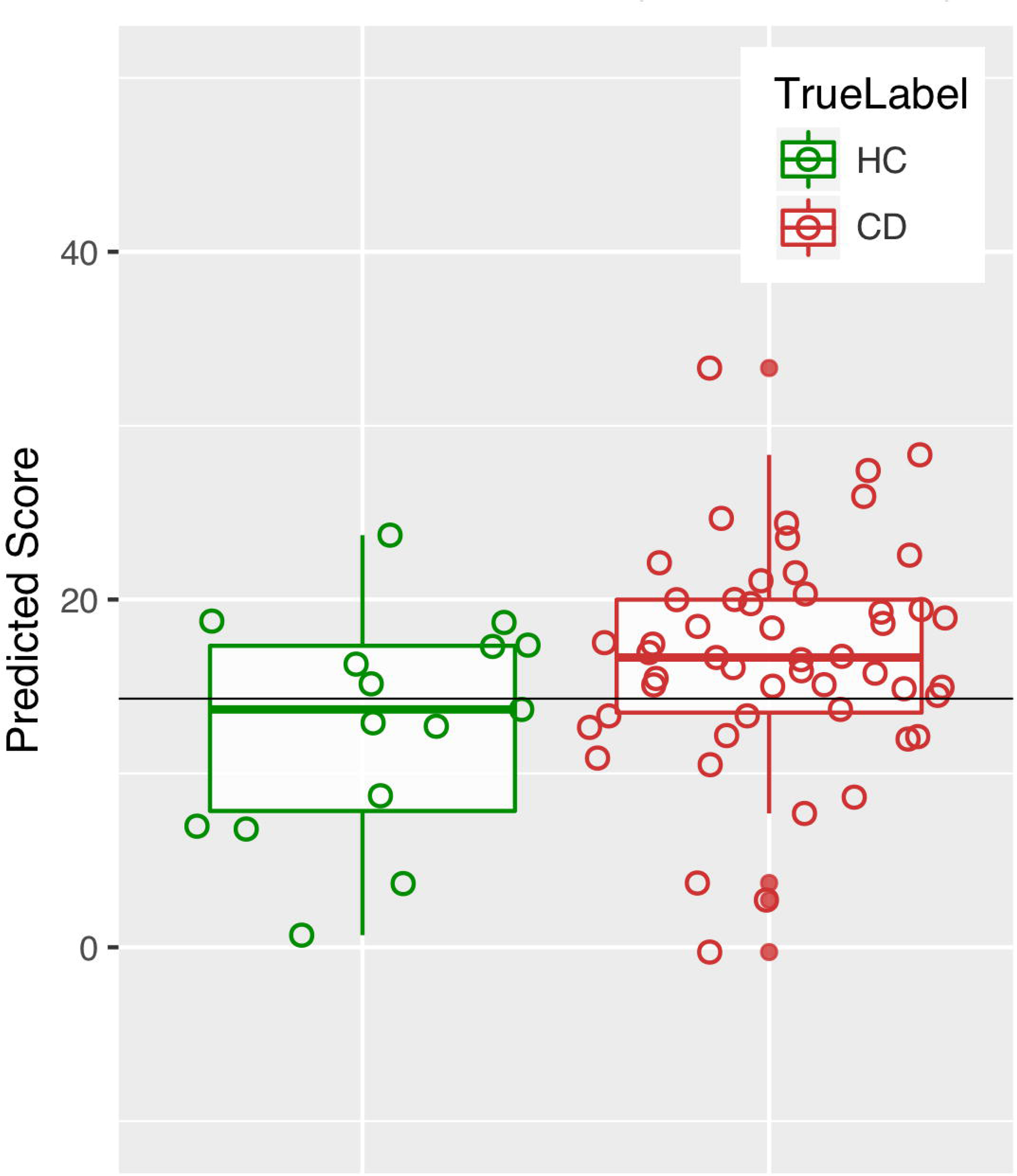
Prediction on CD-test. Predictions are made via SVM models built with *DKMcost*_*125*_ genes as described in text. Batch effect was removed individually for each individual before model building. Black line at x = 14.3 indicates the default cutoff. On average, CD patient scores (red) are higher than those of healthy individuals (green).

## Discussion

After years of study on the subject and numerous promising findings, CD risk prediction from genetic information still remains a problem. We developed AVA,Dx, a machine learning method that uses individual exome data of a panel of Crohn’s disease patients and healthy individuals to select CD-relevant genes and, potentially, predict the health status of previously unseen individuals. We first identified the functional effects[11, 12] of exome SNVs and combined them to create gene scores, indicative of gene functional deficiencies. This approach efficiently decreases the dimensionality of data from considering all exome variants (173,013 variants) to focusing only on affected genes (13,957 genes). Additionally, FS techniques highlight the disease-related genes thus further reducing the dimensionality of data. While our method currently only considers coding variants, the path to integrating other CD-relevant types of variants[54], e.g. splice site, regulatory, *etc.*, into gene scoring is also clear.

The main idea behind AVA,Dx is that disease-causing variation is likely to be functionally detrimental to affected genes/pathway components. To evaluate whether molecular function disruption is an important indicator of gene involvement in disease we tested a number of scoring schemes for evaluating variant effect. As expected, models built with *PascalGWAS* genes performed significantly better than random regardless of the scoring scheme, particularly for the larger CD-train panel. These results indicate both that GWAS (with Pascal filtering) indeed captures CD-association successfully and that the captured signal is likely that of association, not causation. On the other hand, *gene_score*-based models performed significantly better than all other schemes when used with FS gene sets, indicating that severity of the variant functional effect contributes to annotating genes involved in causing disease.

In earlier work, logistic regression on GWAS data from a CD panel of ~13,000 individuals attained a ROC AUC of 0.86[8]. However, using only ~1,300 people, reduced performance quite significantly (ROC AUC = 0.60). The number of individuals in this type of study clearly contributes heavily to the success of CD risk prediction. On the other hand, AVA,Dx reached its peak performance (ROC AUC=0.74) using no non-coding variants and only 111 people – less than a tenth involved in GWAS. Interestingly, the logistic regression model described in Wei et al.[8] (albeit limited by using exonic variation only) was able to correctly identify 46 of 64 patients in our cohort, but misidentified 26 of 47 healthy individuals as affected by Crohn’s Disease. AVA,Dx identified just one more patient correctly, but it did so at significantly higher accuracy – mislabeling eight people fewer than logistic regression analysis.

Large sample size requirements and use of common SNPs only, limit applicability of GWAS. Causative factors of CD remain unidentified by GWAS-based models even as their predictive accuracy improves. For instance, some disease-related genes may not be found simply because they contain variants not covered by the SNP-array or because there were not enough people in the study. As mentioned above, GWAS associations are also often markers of disease, rather than causes. Our FS genes (*e.g. KS5*_*max*_ and *DKMcost*_*125*_), on the other hand, are selected based on the functional changes of the genes, descriptive of separating CD-affected individuals from healthy controls. This approach is not only more precise than GWAS, as illustrated by our model performance (0.74 vs. 0.70 for *DKMcost*_*125*_ and *PascalGWAS*_*175*_ models, respectively), but also informative of pathogenicity pathways (Additional file 12). Interestingly, while FS and external gene set-based models both perform well, the sets did not have much overlap, suggesting that FS identifies previously unknown CD-related genes. We also note that for other complex or rare diseases, where GWAS data is not available or informative, AVA,Dx may work to predict health-status and identify pathogenicity pathways based on even a small number of whole exome sequences.

To summarize, we developed AVA,Dx, a method that uses exome variant-caused gene functional changes to identify disease-related genes and makes to make health status predictions. Our method is able to use less than five percent of the people normally involved in a GWAS study to identify disease-genes and to make fairly accurate CD predictions for previously unseen individuals. Notably, AVA,Dx appears to be fairly robust to differences in panels and in sequencing/filtering methods, making it a potentially clinically useful tool. Furthermore, the CD-genes we identify appear to be relevant to CD, as indicated by the matches of our pathways to known work, and yet significantly different from those highlighted by GWAS. Thus, AVA,Dx presents an orthogonal way for identifying disease-related genes, while avoiding the most severe research limitation – the requirement of a large study panel. Note that while our results indicate that a larger panel would be beneficial for our method’s predictive capacity, they also suggest that significant gains can be achieved by simply doubling the number of people in the study to use less than a tenth of what a GWAS study would require. Looking ahead, it is clear that increasing panel size and including the effects of regulatory variants is likely to improve performance. Moreover, we suggest that the AVA,Dx approach to model building is not limited to Crohn’s Disease, but is rather applicable to a wide spectrum of genetically linked, potentially rare and complex, diseases.

## Methods

### Individuals in the study

Two panels of individuals from Germany were used in this study (Supplementary Online Materials, SOM_Tables 1-2). In records, the CD-train panel included 64 unrelated Crohn’s disease affected individuals (CDs) and 47 unrelated healthy controls (HCs; Additional file 17). Since phenotypic differences between adult and pediatric onset CD suggest different molecular and genetic predisposition pathways[46–49], we further divided CD-train panel into CD-train_early_ and CD-train_late_ panels. CD-train_early_ included 36 early onset CDs (age of onset before 10-year old) and 47 HCs, and CD-train_late_ included 28 late-onset CDs and the same 47 HCs as in CD-train_early_. The CD-test[55] panel included 51 CDs and 15 HCs (Additional file 18), from 28 different pedigrees, including one monozygotic twin pair discordant for CD and eight unrelated heathy controls from a separate panel.

### Exome sequencing and analysis

Samples from both panels were sequenced using Illumina TruSeq Exome Enrichment Kit and the Illumina HiSeq2000 instrument. Reads were mapped to the human genome build hg19. Samples of each panel were called together using Genome Analysis Toolkit (GATK[56] version 3.3-0) Haplotype Caller. Variant calls were restricted to the TruSeq exome target. The VQSR (Variant Quality Score Recalibration) method was employed to identify true polymorphisms in the samples rather than those due to sequencing, alignment, or data processing artifacts.

For each VCF file we ran ANNOVAR[57] to identify all variants mapping to Swiss-Prot[58] proteins. Specifically, we extracted the RefSeq mRNA identifiers from ANNOVAR output and mapped these to Swiss-Prot. Note that if a single variant mapped to more than one protein, all proteins were included into the affected set.

### Individual relationships

To confirm individual relationships, we used hierarchical cluster analysis (from R package SNPRelate[59]) on the individual SNP dissimilarity matrix to draw dendograms. The dendogram of CD-train implied family relationships for individuals S076&S111 and S087&S110 (Additional file 1: Figure 22), which were not initially recorded in the individual information. In the analyses throughout the study, we considered each pair of them as a family and the individuals from one family were left out as one-fold in the cross-validation analysis (see ***FS candidate gene set extraction*** and ***CD model training***). The individual relationships from CD-test were confirmed to include 28 families (Additional file 1: Figure 23).

### Data filtering

We removed all variant calls on the X- and Y- chromosomes, as well as mitochondrial DNA variants. We then filtered the original VCF files with VQSR and retained only the PASS variants. We computed the number of different types of variants in each panel (Additional file 2 to 5) separately for HC and CD individuals and tested the difference between them using the Kolmogorov-Smirnov test[15]. Since we found significant differences between CDs and HCs within the same data panel (Additional file 2 and 3) and across panels (Additional file 15), we further cleaned the data to remove all variants with missing calls (identified as ./.). Removal of these loci ensured that every individual has a confident call at every locus of the same panel. All filtering was done using VCFtools[16].

### Association tests

We applied association tests to the VCF files using PLINK[17]. Specifically, 2-sided Fisher’s Exact test was used to calculate the p-value from the CD-HC allelic counts of each variant. When there were multiple alternative variants, p-values of all possible variants were calculated. Benjamini-Hochberg[18] correction was applied to attained p-values to correct for multiple hypothesis testing.

### Gene scoring

We first checked the Swiss-Prot protein sequence for correspondence; *i.e.* we looked for the variant-defined wild-type residue to exist in the variant sequence position. If the position contained the mutant amino acid instead, we assumed allele disagreement between reference databases RefSeq and Swiss-Prot. For these variants, we chose the RefSeq sequence to be correct and replaced the amino acid in the Swiss-Prot sequence to correspond to RefSeq. We then computed the raw SNAP[11] score for each variant, ranged from -100 to 100, where any score less than or equal to zero is classified as neutral, *i.e.* no protein function change, and non-neutral otherwise.

An individual *variant score* was assigned as follows, for:

1. non-synonymous variants

a. SNAP score >=0 (effect): v_score=0.06+(SNAP score/100)*0.94
b. SNAP <0 (neutral): v_score = 0.055
2. synonymous variant, v_score = 0.05
3. InDel variants, v_score = 1
4. erroneously mapped variants and variants in 11 genes that could not be handled by SNAP (genes > 6000 amino acids), v_score = 0.055.

Individual v_scores of heterozygous variants were multiplied by 0.25 (in Eqn. 1 *het*=0.25 for heterozygous and *het*=1 for homozygous variants) to approximate the effects of heterozygosity.

For every gene in every individual we computed a gene functional deficit score (*gene_score*) as a sum over all gene-specific *v_scores* (Eqn. 1). Note that gene scores computed in this fashion are zero only for genes that have no variants at all. However, further comparison between gene scores for different genes is not possible, as the score is highly dependent on gene length and overall tolerance for variability, *e.g.* longer genes with more variable regions will tend to score higher while remaining relatively functional biochemically.

Thus, for each gene, *g*, the overall variant burden score of all *N*_*g*_ variants was:

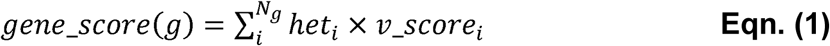

We also calculated other versions of the *gene_score*:

1. an easy_*gene_score*, as in Eqn. 1, but non-synonymous v_score=1
2. a binary_*gene_score*, as in Eqn. 1, but SNAP-neutral non-synonymous v_score=0.055 and effect non-synonymous v_score=1
3. *EV_gene_score*, in which gene_score = # of exonic variants in the gene
4. *NSV_gene_score*, in which gene_score = # of nonsynonymous variants in the gene

In our representation, thus, every individual exome can thus be viewed as a vector of individual gene scores with an associated binary disease class (status: CD *vs.* HC). All exome vectors of one panel of individuals are of the same length; *i.e.* genes that are not affected by any variants in a particular individual are assigned a zero score. Genes with no variants in any individual in a panel were removed from consideration. Additionally, we also removed genes that have only one non-zero score (one individual with a variant) and genes that have consistent non-zero scores across a panel (all have the same variant).

### Reference candidate gene set extraction

The *ALL set* represented all genes where at least one individual in CD-train had at least one variant.

For each human protein entry in Swiss-Prot (20,204 entries, September 13^th^, 2015) set we extracted all gene and protein names, mapped to the protein entry identifier. All single word names longer than two characters were included into the search. The Swiss-Prot identifier, minus the “_HUMAN” suffix was also included as a name. Names were normalized as follows:

1. Multi-word names were converted into bags-of-words. The abstracts were searched for a combination of all words.
2. In a multi-word name, the word separators were any combination of space “ ”, parenthesis “(”, “)”, colon “:”, or semicolon “;”.
3. All single words were normalized by replacing any dash “-“, forward slash “/”, star “*”, comma “,”, carrot “^”, and period “.” with a star “*” in the middle of a word and by removing them at the ends.

All PubMed abstracts were normalized in the same fashion as names. We searched these abstracts for the occurrence of names in CD publications (“crohn disease” MeSH term, *MeSH set*, 2,471 genes, computed on October 27^th^, 2015, and 1824 genes from *MeSH set* were in CD-train, Additional file 19).

In addition to mining the literature, we also used

1. a set of all 1,286 validated protein-coding genes in the 163 known CD-associated regions[3] (*GWAS set*, 925 genes from *GWAS set* were in CD-train, Additional file 7)
2. a Pascal[9] ranked list of CD-related genes, 393 genes with a Benjamini & Hochberg[18] corrected p-val < 0.05 (*PascalGWAS set*, Additional file 8, 318 genes from *PascalGWAS set* were in CD-train)
3. a set of all proteins annotated as Crohn’s disease related (Disease feature) in Swiss-Prot (*SP set*, 22 genes, February 29^th^, 2016, and 18 genes from *SP set* were in CD-train, Additional file 20)

*MeSH*, *GWAS*, *PascalGWAS* and *SP* gene sets were considered *external sets* throughout the study. In text, a subscript number following the set name indicated the gene number of top ranked genes from this set used to build models. For example, *PascalGWAS*_*100*_ indicated building a model using top ranked 100 genes from *PascalGWAS set*. A subscript letter *r* before the gene number meant a random selection of genes from this set instead of top ranked genes (e.g. *GWAS*_*r100*_ indicated 100 genes randomly selected from *GWAS set* were used for model building).

### Features selection (FS) candidate gene set extraction

We recorded genes affected by at least one variant (*ALL set*) separately for each CD-train, CD-train_early,_ and CD-train_late_ panels. We performed the following gene set selections:

1. Collected genes where at least 3 CDs and no HCs had non-zero *gene_scores* (*Disease set*, abbreviated as *DIS set*).
2. Compared the distribution of five different versions, respectively, of *gene_scores* for CDs *vs.* HCs using the *t-test* (*TT5 set*) and *ks-test* (*KS5 set*) and took the genes that were differently (p < 0.05, no correction for multiple testing) distributed in CDs and HCs.
3. Applied DKMcost[60] feature selection (from R CORElearn package[61]) and ranked genes by their merit.

In order to avoid overfitting, we applied the above FS techniques (*DIS*, *KS5*, *TT5* and *DKMcost)* in a *leave one person out* fashion, iteratively in each fold of cross-validation. Thus, we had built multiple models of CD-train-based AVA,Dx with different gene sets each using the same FS technique, so that no model was trained and tested on the same sample. As a “sanity check”, we collected genes as described in method (1) above from the entire CD-train (overfitting), and trained *DISO* gene models in a *leave-one-out* fashion. Subscript numbers, *e.g. KS5*_*r100*_ or *DKMcost*_*125*_, meant the random (_r_) or top ranked, respectively, number of genes used in building the model as described in *Reference candidate gene set extraction.* When all genes from the *FS set* were used to build a model, the gene name was followed by a subscript *max* (*e.g. KS5*_*max*_).

### Computing gene set overlap

As described above, the number of FS candidate gene sets for one panel and one extraction technique was equal to the number of unrelated individuals in that panel, *e.g.* there were 109 different *KS5 sets* in a 109-fold cross-validation on CD-train data. For calculation of overlap between any *KS5 set* and a gene set with fixed genes, *e.g. MeSH set*, we computed the overlap and the significance (hypergeometric distribution test against a background of the corresponding variant-affected genes) for all 109 *KS5 sets* and recorded the mean. For calculation of overlap between two non-fixed gene sets, *e.g.* between *KS5* and *DKMcost sets*, we computed the overlap and significance when the same test individual was held-out, and recorded the mean.

### Finding gene networks

We used the ConsensusPath database[27] to identify the enrichment in alterations of the known molecular pathways in the selected CD-train genes. *ALL set* of CD-train was used as the background list. *KS5*_*max*_ and *DKMcost*_*125*_ selected from entire CD-train panel, as well as *PascalGWAS*_*175*_ genes, were used as input for the pathway enrichment analysis. All pathways with a q-val < 0.1 were recorded in Additional file 12. Induced network analysis from ConsensusPath database using FS genes as starting points was used to detect additional potentially CD-associated genes.

### CD models

#### Training cross-validation

We built CD models using *leave one person out* cross-validation on the CD-train panel. Note that individuals of the pair S087 & S110 and the pair S076 & S111 are more genetically similar than others (and potentially related, Additional file 1: Figure 22), so we left the members of each of these pairs out simultaneously in our *leave-one-out* cross-validation, *i.e.* we performed a 109-fold cross-validation on the CD-train data of 111 individuals. For each model, to make the classes of the training set balanced for healthy vs. CD individuals, we bootstrapped the individual samples of the minor class (resampling with replacement) to create new training samples in a balanced manner. All models used the Support Vector Machine (SVM) algorithm in R’s e1071 package[62]. Note that changing the learning method, *i.e.* replacing support vector machines with Naïve Bayes, neural networks, *etc.* or adjusting method parameters, could potentially produce better results. However, as the goal of this experiment was to evaluate the CD-relevance of the selected gene sets, we did not optimize algorithm performance. For evaluating the performance of the different gene selection methods, we:

1. randomly sampled with replacement different numbers (10, 25-200, in steps of 25, 500, or 1000) of genes, 100 times from each cross-validation fold gene-set. The gene number was recorded as a subscript following the gene set name as described in *Reference candidate gene set extraction* and *FS candidate gene set extraction* sections. For example, for 100-gene *KS5 set* (*KS5r*_*100*_) in CD-train this meant that we trained 10,900 models – 100 random gene sets for each cross-validation fold. Note that when the gene set did not have enough genes for sampling, we used the entire set to build models – one model per fold (*max* subscript following the gene set name; *e.g. KS5_max_*). For example, if the *KS5 set* had 113 genes, models requiring more than that used the whole KS5 (*KS5*_*max*_) set in every model training iteration. Note that for all models built using a fixed set of genes, the only source of difference in model performance is the differential resampling of the training individuals of the minor class.
2. we took the top ranked 10 or 25 to 200 (in steps of 25) genes and performed cross-validation with the same top ranked genes for each fold of cross-validation: *e.g. DKMcost*_*50*_ means we trained 10,900 models – top ranked 50 *DKMcost* genes for each testing fold. Here as above, the only source of the difference in model performance is the differential resampling of the training individuals of the minor class.

For each gene set we computed the various model performance metrics, including area under (AUC) of the precision-recall (PR) and ROC curves (R package PRROC[63], Eqn. 2, where TP = true positives, correctly identified individuals with CD; FP = false positives, healthy individuals misclassified as having CD; TN = true negatives, correctly identified healthy individuals; FN = individuals with CD misidentified as being healthy).

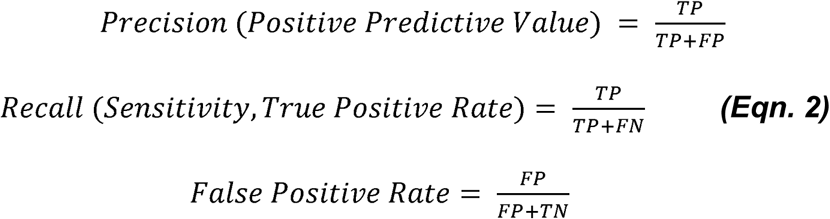

Class labels for CD and HC were set at 100 and -100; *i.e.* more negative scores indicate likely healthy individuals and more positive scores indicate likely CD patients. We obtained ROC and PR curves for by varying the threshold for classifying an individual as CD-affected or healthy from -100 (most healthy) to 100 (most CD).

To further test if the performance was achieved by chance, we did permutation test by shuffling the labels of every training fold in the cross-validation 1,000 times. Null distributions of ROC and PR AUCs were based on these 1,000 results. The permutation p-values for the “real” cross-validation AUCs were obtained empirically by counting the number of larger permuted AUCs (divided by 1000).

#### Logistic regression from GWAS data

We obtained the logistic regression from a previous study[8] via personal contact. CD-train has 31 loci among the 573-loci regression model, and we made prediction with the model using the 31 loci (listed in Additional file 21). We obtained predictions of precision=64% and recall=72% with this model.

#### Prediction: Eliminating inter-panel batch effects of sequencing

To remove the batch effect, we applied the ComBat[53] method (from R package sva[64]). ComBat was developed originally for gene expression data. It is an empirical Bayes framework for adjusting data for batch effect[53]. Here we apply the parametric ComBat to the *gene_score,* which represents the gene functional change instead of expression change.

We first applied ComBat to the combined panel of CD-train and CD-test. We checked, using *pam*[52] (R package cluster[65]) if individuals in this combined set can be clustered correctly into the original batches (CD-train and CD-test). Jaccard coefficient was used to evaluate the cluster similarity between the original batches and clustering results. We then performed the training cross-validation (as described in *Training cross-validation*) with ComBat-adjusted *gene_score* for the *DKMcost*_*125*_ gene set to confirm that the adjustment did not alter the signal in the data. To simulate the “real-world situation” of predicting disease, ComBat was applied individually to each person of CD-test in combination with the entire CD-train panel. Note that since in this case the unknown batch has only one sample, only the means, and not the variance, of the *gene_scores* were adjusted for batch effect. In all cases, as ComBat requires a class label (here, health status) of every input individual, we assigned 0 (unknown) as the health status for each test individual, where 100 indicated CD and -100 indicated HC in the CD-train panel.

#### Prediction: Model-Building

We used the entire unadjusted CD-train panel to select *DKMcost*_*125*_ *genes* as fixed features for our final model. We then tested the predictive ability of our model by predicting the health status of 66 individuals from CD-test panel. Each individual was predicted separately, as described above, to mimic the real-world situation. After removal of batch effect, we resampled the CD-train panel to create 500 individuals of each HC and CD class and used the model to predict on the individual.

#### Prediction: Choosing the default cutoff

We once more built models of CD-train in cross-validation as described above, except that this time we also resampled individuals in each fold of training to create 500 individuals of the CD and HC classes. We used the original selected *DKMcost*_*125*_ gene sets for each of the 109 models and tested on the left-out individuals. We computed the means of the prediction scores of CD individuals and of HCs and chose the mean of these two means as the default cutoff. As the cutoff varied with different resampling and training rounds, we conducted this process 100 times and chose the most common cutoff value (Additional file 1: Figure 20) for subsequent predictions. When predictions of all individuals were made, we evaluated the performance by ROC/PR AUCs and the MCC (Matthews correlation coefficient[66], Eqn. 3) based on the chosen cutoff. The significance of the prediction result was also evaluated by a 1,000-time permutation test as described in *cross-validation,* except that we shuffled the labels for the entire CD-train panel this time and null distributions of ROC and PR AUCs were drawn from the prediction results on individuals from CD-test panel.

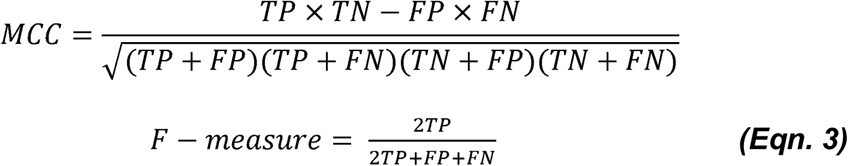

## List of abbreviations

IBD: Inflammatory bowel disease
CD: Crohn’s disease
HC: Healthy control
GWAS: Genome wide association study AVA
Dx: Analysis of Variation for Association with Disease
ROC: Receiver operating characteristic
PR: Precision Recall
AUC: Area under the curve
FS: Feature selection

## Declarations

### Ethics approval and consent to participate

Patient recruitment was approved by the Ethics Committee of the University Hospital S.-H. In Kiel (ID A156/03), and each patient gave their informed consent.

### Consent for publication

Not applicable.

### Availability of data and material

VCF data of the training and testing panels are accessible at the Critical Assessment of Genome Interpretation website (CAGI, https://genomeinterpretation.org/content/4-crohns-exomes).

### Competing interests

The authors declare that they have no competing interests.

### Funding

YB and YW were supported by the NIH U01 GM115486 and the Informatics Research Starter grant from the PhRMA foundation. YB was also supported by U24 MH06845. This study was supported by the German Ministry of Education and Research (BMBF) program e:Med sysINFLAME (http://www.gesundheitsforschung-bmbf.de/de/5111.php, no.: 01ZX1306A) and received infrastructure support from the Deutsche Forschungsgemeinschaft (DFG) Cluster of Excellence ‘Inflammation at Interfaces’ (http://www.inflammation-at-interfaces.de, no.: XC306/2).

### Authors' contributions

Y.A. developed code for gene scoring and processing of PubMed abstracts. B.P. and A.F. provided the CD data. Y.W. developed and optimized the gene selection methods and conducted all analyses; Y.B. conceived the methods and supervised all work. Y.B. and Y.W. wrote the manuscript. All authors participated in manuscript revisions and improvement.

## Acknowledgements

We would like to thank Dr. Jay Tischfield, Dr. Derek Gordon, Dr. Jinchuan Xing (all Rutgers) for all help and comments that greatly improved the manuscript. We are also grateful to Dr. Burkhard Rost (TU Munich), Dr. Predrag Radivojac (U Indiana), Dr. Vikas Nanda, Yannick Mahlich, Max Miller, Dr. Chengsheng Zhu, and Dr. Anton Molyboha (all Rutgers) for all discussions.

We would also like to express gratitude to all people of this study who have made their genomic and medical information available to contribute to a better understanding of this disease.

## REFERENCES

1. Lakatos PL: Recent trends in the epidemiology of inflammatory bowel diseases: up or down? World J Gastroenterol 2006, 12:6102–6108.

2. Chen GB, Lee SH, Brion MJ, Montgomery GW, Wray NR, Radford-Smith GL, Visscher PM: Estimation and partitioning of (co)heritability of inflammatory bowel disease from GWAS and immunochip data. Hum Mol Genet 2014, 23:4710–4720.

3. Jostins L, Ripke S, Weersma RK, Duerr RH, McGovern DP, Hui KY, Lee JC, Schumm LP, Sharma Y, Anderson CA, et al: Host-microbe interactions have shaped the genetic architecture of inflammatory bowel disease. Nature 2012, 491:119–124.

4. Silverberg M, Daly M, Moskovitz D, Rioux J, McLeod R, Cohen Z, Greenberg G, Hudson T, Siminovitch K, Steinhart A: Diagnostic misclassification reduces the ability to detect linkage in inflammatory bowel disease genetic studies. Gut 2001, 49:773–776.

5. Henriksen M, Jahnsen J, Lygren I, Sauar J, Schulz T, Stray N, Vatn MH, Moum B, Group IS: Change of diagnosis during the first five years after onset of inflammatory bowel disease: results of a prospective follow-up study (the IBSEN Study). Scandinavian journal of gastroenterology 2006, 41:1037–1043.

6. Ruderfer DM, Korn J, Purcell SM: Family-based genetic risk prediction of multifactorial disease. Genome Med 2010, 2:2.

7. Jostins L, Barrett JC: Genetic risk prediction in complex disease. Hum Mol Genet 2011, 20:R182–188.

8. Wei Z, Wang W, Bradfield J, Li J, Cardinale C, Frackelton E, Kim C, Mentch F, Van Steen K, Visscher PM: Large sample size, wide variant spectrum, and advanced machine-learning technique boost risk prediction for inflammatory bowel disease. The American Journal of Human Genetics 2013, 92:1008–1012.

9. Lamparter D, Marbach D, Rueedi R, Kutalik Z, Bergmann S: Fast and Rigorous Computation of Gene and Pathway Scores from SNP-Based Summary Statistics. PLoS Comput Biol 2016, 12:e1004714.

10. Samuels DC, Han L, Li J, Quanghu S, Clark TA, Shyr Y, Guo Y: Finding the lost treasures in exome sequencing data. Trends Genet 2013, 29:593–599.

11. Bromberg Y, Rost B: SNAP: predict effect of non-synonymous polymorphisms on function. Nucleic acids research 2007, 35:3823–3835.

12. Hecht M, Bromberg Y, Rost B: Better prediction of functional effects for sequence variants. BMC genomics 2015, 16:S1.

13. Chang C-C, Lin C-J: LIBSVM: a library for support vector machines. ACM transactions on intelligent systems and technology (TIST) 2011, 2:27.

14. DePristo MA, Banks E, Poplin R, Garimella KV, Maguire JR, Hartl C, Philippakis AA, Del Angel G, Rivas MA, Hanna M: A framework for variation discovery and genotyping using next-generation DNA sequencing data. Nature genetics 2011, 43:491–498.

15. Massey Jr FJ: The Kolmogorov-Smirnov test for goodness of fit. Journal of the American statistical Association 1951, 46:68–78.

16. Danecek P, Auton A, Abecasis G, Albers CA, Banks E, DePristo MA, Handsaker RE, Lunter G, Marth GT, Sherry ST, et al: The variant call format and VCFtools. Bioinformatics 2011, 27:2156–2158.

17. Purcell S, Neale B, Todd-Brown K, Thomas L, Ferreira MA, Bender D, Maller J, Sklar P, De Bakker PI, Daly MJ: PLINK: a tool set for whole-genome association and population-based linkage analyses. The American Journal of Human Genetics 2007, 81:559–575.

18. Benjamini Y, Hochberg Y: Controlling the false discovery rate: a practical and powerful approach to multiple testing. Journal of the royal statistical society Series B (Methodological) 1995:289–300.

19. Lek M, Karczewski KJ, Minikel EV, Samocha KE, Banks E, Fennell T, O'Donnell-Luria AH, Ware JS, Hill AJ, Cummings BB, et al: Analysis of protein-coding genetic variation in 60,706 humans. Nature 2016, 536:285–291.

20. Johnson NL, Kemp AW, Kotz S: Univariate discrete distributions. John Wiley & Sons; 2005.

21. Liu Z, Lenardo MJ: The role of LRRK2 in inflammatory bowel disease. Cell Res 2012, 22:1092–1094.

22. Yamamoto S, Ma X: Role of Nod2 in the development of Crohn's disease. Microbes Infect 2009, 11:912–918.

23. Cho JH, Brant SR: Recent insights into the genetics of inflammatory bowel disease. Gastroenterology 2011, 140:1704–1712.

24. Skovdahl HK, Granlund A, Ostvik AE, Bruland T, Bakke I, Torp SH, Damas JK, Sandvik AK: Expression of CCL20 and Its Corresponding Receptor CCR6 Is Enhanced in Active Inflammatory Bowel Disease, and TLR3 Mediates CCL20 Expression in Colonic Epithelial Cells. PLoS One 2015, 10:e0141710.

25. Canto E, Garcia Planella E, Zamora-Atenza C, Nieto JC, Gordillo J, Ortiz MA, Meton I, Serrano E, Vegas E, Garcia-Bosch O, et al: Interleukin-19 impairment in active Crohn's disease patients. PLoS One 2014, 9:e93910.

26. Bretin A, Carriere J, Dalmasso G, Bergougnoux A, B'Chir W, Maurin AC, Muller S, Seibold F, Barnich N, Bruhat A, et al: Activation of the EIF2AK4-EIF2A/eIF2alpha-ATF4 pathway triggers autophagy response to Crohn disease-associated adherent-invasive Escherichia coli infection. Autophagy 2016, 12:770–783.

27. Kamburov A, Stelzl U, Lehrach H, Herwig R: The ConsensusPathDB interaction database: 2013 update. Nucleic Acids Res 2013, 41:D793–800.

28. Wehkamp J, Schmid M, Stange EF: Defensins and other antimicrobial peptides in inflammatory bowel disease. Current opinion in gastroenterology 2007, 23:370–378.

29. Nuding S, Fellermann K, Wehkamp J, Stange EF: Reduced mucosal antimicrobial activity in Crohn's disease of the colon. Gut 2007, 56:1240–1247.

30. Guaní-Guerra E, Santos-Mendoza T, Lugo-Reyes SO, Terán LM: Antimicrobial peptides: general overview and clinical implications in human health and disease. Clinical Immunology 2010, 135:1–11.

31. Bevins CL, Salzman NH: Paneth cells, antimicrobial peptides and maintenance of intestinal homeostasis. Nature Reviews Microbiology 2011, 9:356–368.

32. Kim EK, Choi E-J: Pathological roles of MAPK signaling pathways in human diseases. Biochimica et Biophysica Acta (BBA)-Molecular Basis of Disease 2010, 1802:396–405.

33. Mudter J, Neurath MF: Apoptosis of T cells and the control of inflammatory bowel disease: therapeutic implications. Gut 2007, 56:293–303.

34. Claret L, Miquel S, Vieille N, Ryjenkov DA, Gomelsky M, Darfeuille-Michaud A: The flagellar sigma factor FliA regulates adhesion and invasion of Crohn disease-associated Escherichia coli via a cyclic dimeric GMP-dependent pathway. Journal of Biological Chemistry 2007, 282:33275–33283.

35. Khoshakhlagh P, Bahrololoumi-Shapourabadi M, Mohammadirad A, Ashtaral-Nakhai L, Minaie B, Abdollahi M: Beneficial effect of phosphodiesterase-5 inhibitor in experimental inflammatory bowel disease; molecular evidence for involvement of oxidative stress. Toxicology mechanisms and methods 2007, 17:281–288.

36. Kolios G, Valatas V, Ward SG: Nitric oxide in inflammatory bowel disease: a universal messenger in an unsolved puzzle. Immunology 2004, 113:427–437.

37. Hällgren R, Colombel JF, Dahl R, Fredens K, Kruse A, Jacobsen NO, Venge P, Rambaud JC: Neutrophil and eosinophil involvement of the small bowel in patients with celiac disease and Crohn's disease: studies on the secretion rate and immunohistochemical localization of granulocyte granule constituents. The American journal of medicine 1989, 86:56–64.

38. Witko-Sarsat V, Pederzoli-Ribeil M, Hirsh E, Sozzani S, Cassatella MA: Regulating neutrophil apoptosis: new players enter the game. Trends in immunology 2011, 32:117–124.

39. Dai C, Jiang M, Sun M-J: Innate immunity and adaptive immunity in Crohn's disease. Annals of translational medicine 2015, 3.

40. Korzenik J: The Role of Innate Immunity in Crohn's Disease. Gastroenterology & hepatology 2007, 3:82.

41. Marcil V, Sinnett D, Seidman E, Boudreau F, Gendron F-P, Beaulieu J-F, Menard D, Lambert M, Bitton A, Sanchez R: Association between genetic variants in the HNF4A gene and childhood-onset Crohn's disease. Genes and immunity 2012, 13:556.

42. Consortium UIG, 2 WTCCC: Genome-wide association study of ulcerative colitis identifies three new susceptibility loci, including the HNF4A region. Nature genetics 2009, 41:1330.

43. Babeu J-P, Boudreau F: Hepatocyte nuclear factor 4-alpha involvement in liver and intestinal inflammatory networks. World journal of gastroenterology: WJG 2014, 20:22.

44. Belkina AC, Denis GV: BET domain co-regulators in obesity, inflammation and cancer. Nature reviews Cancer 2012, 12:465.

45. Denis GV: Bromodomain coactivators in cancer, obesity, type 2 diabetes, and inflammation. Discovery medicine 2010, 10:489.

46. Guariso G, Gasparetto M, Visona Dalla Pozza L, D'Inca R, Zancan L, Sturniolo G, Brotto F, Facchin P: Inflammatory bowel disease developing in paediatric and adult age. J Pediatr Gastroenterol Nutr 2010, 51:698–707.

47. Nieuwenhuis EE, Escher JC: Early onset IBD: what's the difference? Dig Liver Dis 2008, 40:12–15.

48. Coughlan A, Wylde R, Lafferty L, Quinn S, Broderick A, Bourke B, Hussey S: A rising incidence and poorer male outcomes characterise early onset paediatric inflammatory bowel disease. Aliment Pharmacol Ther 2017, 45:1534–1541.

49. Michail S, Bultron G, Depaolo RW: Genetic variants associated with Crohn's disease. Appl Clin Genet 2013, 6:25–32.

50. Warr A, Robert C, Hume D, Archibald A, Deeb N, Watson M: Exome sequencing: current and future perspectives. G3: Genes| Genomes| Genetics 2015, 5:1543–1550.

51. Parla JS, Iossifov I, Grabill I, Spector MS, Kramer M, McCombie WR: A comparative analysis of exome capture. Genome biology 2011, 12:1.

52. Reynolds AP, Richards G, de la Iglesia B, Rayward-Smith VJ: Clustering rules: a comparison of partitioning and hierarchical clustering algorithms. Journal of Mathematical Modelling and Algorithms 2006, 5:475–504.

53. Johnson WE, Li C, Rabinovic A: Adjusting batch effects in microarray expression data using empirical Bayes methods. Biostatistics 2007, 8:118–127.

54. Pal LR, Moult J: Genetic Basis of Common Human Disease: Insight into the Role of Missense SNPs from Genome-Wide Association Studies. J Mol Biol 2015, 427:2271–2289.

55. Ellinghaus D, Zhang H, Zeissig S, Lipinski S, Till A, Jiang T, Stade B, Bromberg Y, Ellinghaus E, Keller A, et al: Association between variants of PRDM1 and NDP52 and Crohn's disease, based on exome sequencing and functional studies. Gastroenterology 2013, 145:339–347.

56. McKenna A, Hanna M, Banks E, Sivachenko A, Cibulskis K, Kernytsky A, Garimella K, Altshuler D, Gabriel S, Daly M, DePristo MA: The Genome Analysis Toolkit: a MapReduce framework for analyzing next-generation DNA sequencing data. Genome Res 2010, 20:1297–1303.

57. Wang K, Li M, Hakonarson H: ANNOVAR: functional annotation of genetic variants from high-throughput sequencing data. Nucleic acids research 2010, 38:e164–e164.

58. Consortium TU: UniProt: the universal protein knowledgebase. Nucleic Acids Res 2017, 45:D158–d169.

59. Zheng X, Levine D, Shen J, Gogarten SM, Laurie C, Weir BS: A high-performance computing toolset for relatedness and principal component analysis of SNP data. Bioinformatics 2012, 28:3326–3328.

60. Dietterich T, Kearns M, Mansour Y: Applying the weak learning framework to understand and improve C4. 5. In ICML. 1996: 96–104.

61. Robnik M: Package ‘CORElearn‘. 2015.

62. Meyer D, Dimitriadou E, Hornik K, Weingessel A, Leisch F: e1071: Misc Functions of the Department of Statistics, Probability Theory Group (Formerly: E1071), TU Wien, 2015. R package version:1.6-7.

63. Grau J, Grosse I, Keilwagen J: PRROC: computing and visualizing precision-recall and receiver operating characteristic curves in R. Bioinformatics 2015, 31:2595–2597.

64. Leek JT, Johnson WE, Parker HS, Jaffe AE, Storey JD: The sva package for removing batch effects and other unwanted variation in high-throughput experiments. Bioinformatics 2012, 28:882–883.

65. Maechler M, Rousseeuw P, Struyf A, Hubert M, Hornik K: Cluster: cluster analysis basics and extensions. R package version 2012, 1:56.

66. Matthews BW: Comparison of the predicted and observed secondary structure of T4 phage lysozyme. Biochimica et Biophysica Acta (BBA)-Protein Structure 1975, 405:442–451.

